# CrossTalkeR: Analysis and Visualisation of Ligand Receptor Networks

**DOI:** 10.1101/2021.01.20.427390

**Authors:** James S. Nagai, Nils B. Leimkühler, Michael T. Schaub, Rebekka K. Schneider, Ivan G. Costa

## Abstract

**Motivation:** Ligand-receptor (LR) network analysis allows the characterization of cellular crosstalk based on single cell RNA-seq data. However, current methods typically provide a list of inferred LR interactions and do not allow the researcher to focus on specific cell types, ligands or receptors. In addition, most of these methods cannot quantify changes in crosstalk between two biological phenotypes.

**Results:** CrossTalkeR is a framework for network analysis and visualisation of LR interactions. CrossTalkeR identifies relevant ligands, receptors and cell types contributing to changes in cell communication when contrasting two biological phenotypes, i.e. disease vs. homeostasis. A case study on scRNA-seq of human myeloproliferative neoplasms reinforces the strengths of CrossTalkeR for characterisation of changes in cellular crosstalk in disease.

**Availability and Implementation:** CrosstalkeR is an R package available at https://github.com/CostaLab/CrossTalkeR.

## 1 Introduction

Understanding cellular crosstalk is vital for uncovering molecular mechanisms associated to cell differentiation and disease progression. Single cell RNA sequencing (scRNA-seq) enables the characterization of tissue heterogeneity at an unprecedented level^1, 2^. However, information of cellular proximity and crosstalk is not directly captured by common single cell sequencing protocols. Computational methods, which search for pairs of cell types expressing compatible ligand-receptor (LR) pairs, have thus become a powerful approach for dissecting cellular crosstalk from scRNA-seq data^2^. Current state-of-art LR inference methods, such as CellphoneDB (CPDB)^3^, basically provide a ranked list of hundreds of ligand-receptor and cell pairs for a given scRNA-seq dataset. As interpreting such ranked lists is challenging, this calls for tools to simplify the complexity of these results, i.e., ranking of cell types or genes within a pair of cell types. Also, most LR inference methods focus on the analysis of a single phenotype scRNA-seq experiment and are not able to characterize changes in cellular crosstalk between pairs of phenotypes, e.g., disease vs. homeostasis.

To fill these gaps, we developed CrossTalkeR, which aggregates predictions from CPDB^3^ as cell and cell/gene networks. First it ranks cell types, cell-receptors and cell-ligand pairs regarding their importance in cellular crosstalk based on certain network topology measures. Second, CrossTalkeR enables researchers to perform a phenotype based comparative LR analysis. CrossTalkeR thus complements available methods being the only approach exploring LR networks at both cell and cell-gene levels and by contrasting these networks at pairs of phenotypes (see Supp. Material for an overview of competing methods). CrossTalkeR is implemented as an R package. It presents the results either graphically or in tabular form summarized as a HTML report/PDF report and stores all results in an R container to foster data sharing and reproducibility of results.

## 2 Overview and implementation

The input to CrossTalkeR is a data-table, in which each line has five columns consisting of i) a ligand, ii) a receptor, iii) a sender cell type that expresses the ligand, iv) a receiver cell type that expresses the receptor, and v) a positive score associated with the weight of the interaction, e.g., as predicted by CPDB^3^. Specifically, for CPDB this positive score (weight) corresponds to a function of the mean expression levels of the ligand and the receptor within the respective cell types^3^. From this input data, CrossTalkeR builds network representations and estimates node statistics to highlight particular cells or cell/gene pairs. All the analyses can be performed either on data corresponding to a single phenotype, or data from two phenotypes can be compared.

### 2.1 Network Construction

CrossTalkeR constructs two representations of the LR network: a cell-cell interaction (CCI) network and a cell-gene interaction (CGI) network. In the CCI network, the nodes are defined by each cell-type and the directed edges are weighted by characteristics of the interactions, e.g., number of LR pairs and sum of weights of LR pairs. In the CGI network, the nodes represent gene and cell pairs (ligand with sender cell type, or receptor with receiving cell type) and the edge weights are given by the mean LR expression levels. Edges are always directed from ligands to receptors.

### 2.2 Network topological measures

We use graph theoretic tools to rank nodes (cell types or cell-gene pairs of the CCI (and CGI) network. We compute the following scores (see supplementary material for details):

- *Influencer score*. We compute the number of out-going edges, i.e., the number of signals sent by a cell.
- *Listener score*. We compute the number of in-going edges per node. Nodes with high listener score represent nodes that receive many signals under the studied biological condition.
- *Moderator score*. We compute the betweeness centrality of a node to measure the importance of a node to mediate communication between cells.
- *Importance score*. We characterize the overall *importance* of nodes within the network by computing their *PageRank* score. Note that the *PageRank* can be interpreted probabilistically as the stationary probability distribution of a certain diffusion process on the network.

When comparing two different phenotypes, we obtain differential scores as follows. For the influencer, listener, and moderator scores we simply compute the score difference for each node. For importance (pagerank), we compute log-odds of posteriors probabilities estimated via the pagerank scores.

#### HTML/PDF report and graphical representations

CrossTalkeR creates graphical and tabular representations in the form of a HTML/PDF report. This includes representations of the CCI networks, where edge thickness, color and node sizes represent their characteristics (Fig. 1B). CrossTalker generates PCA plots on the node feature space to describe GCI nodes. It highlights nodes with deviating PCA scores, i.e. nodes outside the 95% confidence interval assuming PCA values are normally distributed (Fig. 1C). Crosstalker also provides a pathway characterization (KEGG pathway enrichment analysis) of top ligand/receptors per node feature to support functional interpretation of results(Fig. 1D). Moreover, it is possible to list all interactions associated within a particular ligand or receptor via a Sankey plot (Fig. 1E). The report also includes tabular representation of rankings, which can be sorted by distinct criteria or searched for particular genes and PC analysis. All results can be exported to spreadsheet formats. The package also produces an R object, which can be used for generation of further graphs, i.e. generation of Sankey plots of ligand or receptors of interest or simplification of plots by filtering ligand, receptors or cells. The software is provided as an open source package (https://github.com/CostaLab/CrossTalkeR). This includes a tutorial to integrate the tool with ligand-receptor analysis tools such as CPDB, as well as example code to reproduce the analysis of stromal cells in human myelofibrosis^4^.

**Figure 1.**
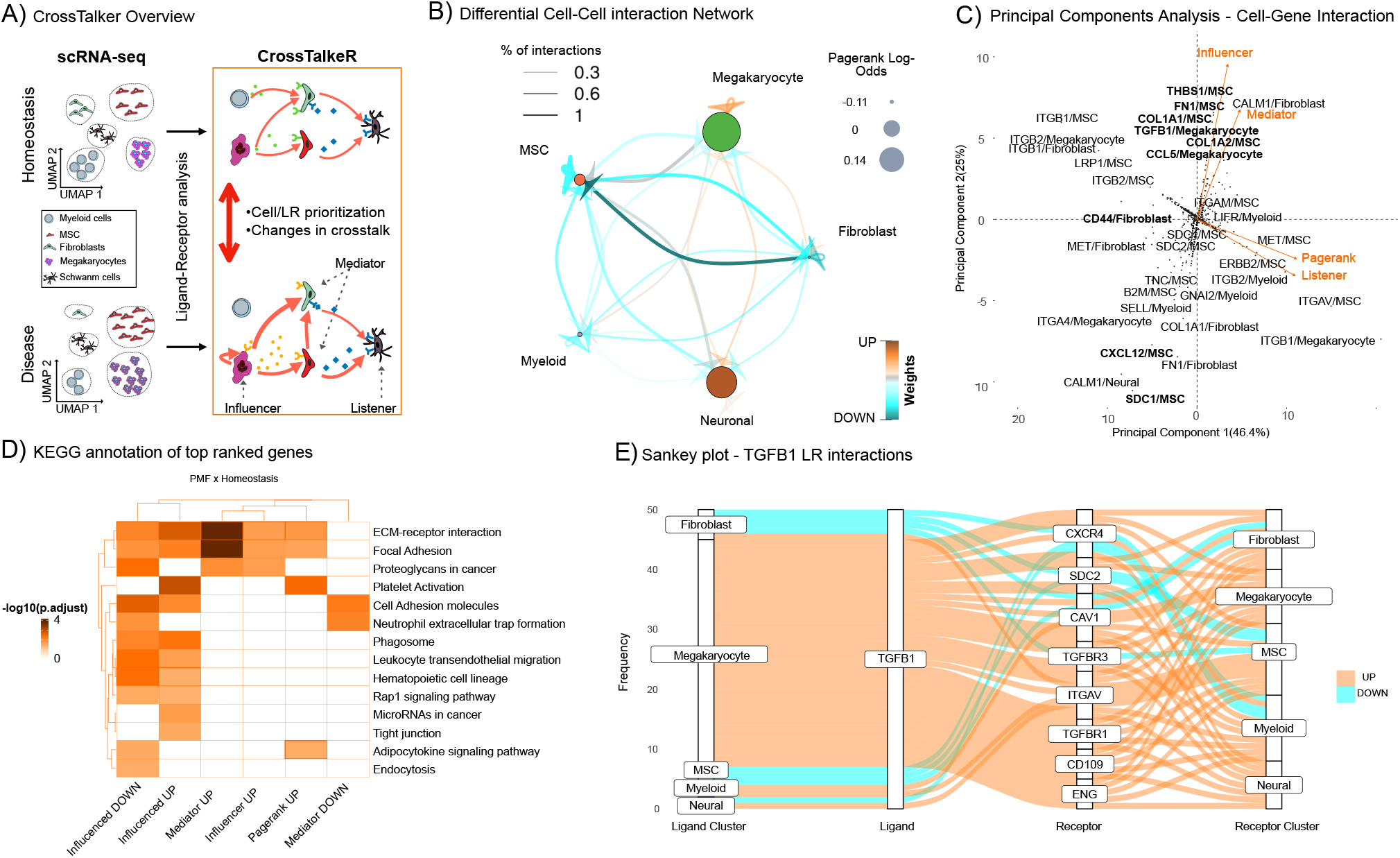
A) CrossTalkeR scheme: given a pair of scRNA-seq, CrossTalkeR creates cell-cell networks (or cell-gene networks), where edge weights are proportional to the expression of ligand-receptors driving communication between the cells. Network topology measure are used to find nodes sending signals (influencers), nodes receiving signals (listener) or both (mediators). Moreover, random-walks estimates with pagerank indicate the importance of network nodes. These measures can be computed for each phenotype individually or in the comparative analyses. B) Comparative Cell-Cell Interaction (CCI), where node size (edge thickness) represents importance of the cell (communication between cells). C) PCA representation of nodes from the Cell Gene Interaction (CGI) based on comparative topological measures. We highlight cell-gene pairs with deviating PCA scores. Only directions associated to positive values are shown (negative values are in opposing order). D) KEGG pathway enrichment provide further insights of genes associated to distinct topological measures (influencer, listener, mediator and node importance) with increase/decrease in disease vs. control (up/down). E) Sankey plot listing all predicted source, receptor and receiver interactions associated with a gene of interest, i.e. TGFB1

## 3 Case study: Cellular crosstalk in myelofibrosis

Here we revisit the scRNAseq analysis comparing bone marrow stromal cells in myelofibrosis (MF) to those in homeostasis^4^. In the comparative CCI network graph (Fig. 1B) megakaryocytes (MK) show a high number of up-regulated autocrine and intercellular interactions. This results in a high pagerank log-odds for this cell population clearly highlighting MK as the most important signaling cells in this disease influencing a plethora of bone marrow cells such as neural cells and MSCs. We also observe a significant loss of interactions involving MSCs and fibroblasts compared to homeostasis. This is in line with the restricted myofibroblastic phenotype MSCs have been described to acquire. MSCs are usually strongly interconnected, moderating the quiescence and differentiation of hematopoietic cells, but loose this capacity as myelofibrosis progresses^4,5^. PCA analysis of the CGI network (Fig. 1C) highlights matrix related proteins (*FN1*, *THBS1*, *COL1A1*), which are secreted by MSCs, and TGFB1, which is secreted by MK, to act as both influencers and mediators in myelofibrosis. These results are in accordance with known mechanisms associated to MF, e.g. the crosstalk between MK and MSCs via TGFb signalling; and the deposition on matrix via MSC cells^6^. Exploration of TGFB1-mediated LR-interactions in the dataset as provided by CrossTalkeR also gives into the differential intercellular signaling present in the myelofibrotic bone marrow (Fig.1E). The strong upregulation of TGFB1-TGFBR1 between MK and MSCs is in line with other publications, showing canonical TGFB1-signaling as most dominant pro-fibrotic element^7^. At the same time, differential expression of co-receptors *CD109* and *ENG* show, how TGFB1-signals are modulated in MF^8,9^. Thus, CrossTalkeR manages to capture the multiplicity of signals expected to be present in biological networks at any given point in time. Similarly, a downregulation of the interaction between *TGFB1* and *TGFRB3*, an endogeneous decoy receptor preventing downstream signaling, also indicates an additional way how TGFB1-signals are increased in MF^10^. Similarly, CXCL12/MSC, a pivotal mediator of hematopoietic homing, is the node with the strongest decrease in the influencer score in disease. This is in line with the findings of the CCI and previously published data showing an extensive loss of hematopoiesis supporting capacities in myelofibrotic MSCs^6^. KEGG pathway analysis (Fig.1D) supports these interpretations with up-regulated mediators and influencers mostly being enriched for ECM^1^ deposition, while down-regulated influenced nodes are associated with hematopoiesis and leukocyte migration. LR analysis with CrossTalkeR also revealed that genes with increased importance are enriched for the KEGG pathway “platelet activation” (Fig. 1D). This recapitulates the finding that platelet related mediators such as *CXCL4* contribute to the perpetual inflammatory environment in myelofibrosis^11^. Taken together, these examples comprehensively illustrate how CrossTalkeR can characterize changes in the diverse cellular crosstalk between conditions from scRNAseq data.

## 4 Conclusion and Future Work

CrossTalkeR is a comprehensive framework for network analysis of LR interactions, and enables comparisons between LR interactions of two biological phenotypes. CrossTalkeR provides both graphical report, tabular data and R objects enabling both interpretable and reproducible analysis. Future work includes the extension of network-based method analysis as well as the analysis of more complex experimental designs as time series or the comparison of several biological conditions.

## Footnotes

1 ECM: Extra Cellular Matrix

