## Supplemental Document 1 for "CrossTalkeR: Analysis and Visualisation of Ligand Receptor Networks"

### Supplementary Material

#### Network Construction

##### *Input Data*

The input to CrossTalkeR consists of a list of 5-tuples $X = \{x_1, ..., x_n\}$ comprising $n$ significant LR interactions predicted by a LR inference tool (e.g., CellPhoneDB) from a single cell RNA-seq data set. Here each 5-tuple is of the form $x = (s, g_{lig}, r, g_{rec}, w)$, where $s \in \{1, .., k\}$ is the sender cell, $r \in \{1, .., k\}$ is the receiver cell, $g_{lig}$ represents a ligand gene, $g_{rec}$ is a receptor gene and $w > 0$ is the weight of the interaction. The Cell types ($s$ and $r$) are defined as one of the $k$ clusters detected in a given scRNA-seq data. The weight $w$ of the interaction indicates the strength in the ligand-receptor signal, e.g., as estimated by the mean expression of the ligand and receptor (MeanLR) in CellPhoneDB.

##### *Cell Gene Interaction based Network*

The vertices $V_{CGI}$ of the Cell-Gene interaction network consist of all possible pairs of ligands with sender cells $(s, g_{lig})$, and receptors with receiver cells $(r, g_{rec})$. Each tuple $(s, g_{lig}, r, g_{rec}, w) \in X$, now gives rise to one directed edge in a weighted graph with vertices $V_{CGI}$. Specifically, the entries of weighted adjacency matrix $\mathbf{W_{CGI}} = \{w_{i,j}\}^{n \times n}$ are defined as $w_{(g_{lig},s),(g_{rec},r)} = w$.

##### *Cell Cell Interaction (CCI) based Network*

We obtain a CCI network by converting the list of 5-tuples $X$ into a list of triplets. Each of these triplets is of the form $t = (s, r, w_{s,r})$, where $s$ represents the sender cell type, $r$ the receiver cell type and $w_{s,r}$ represents the weights of the interactions from cell type $s$ to cell type $r$. Let $W_{s,r}$ be the set of all weights, where $s$ is the sender and $r$ is the receiver cell,

$$W_{s,r} = \{w | (s', g'_{lig}, r', g'_{rec}, w) \in X, s' = s, r' = r\}. \tag{1}$$

Based on these sets, we can define a new weighted directed graph between cells. The vertex set of this graph $V_{CCI}$ consists of the set of all cell types $\{1, .., k\}$, and we denote its adjacency matrix as $\mathbf{W}' = \{w'_{s,r}\}^{k \times k}$, with entries defined as:

$$w'_{s,r} = \sum_{w \in W_{s,r}} w \tag{2}$$

##### *Comparative CGI and CCI Networks*

Given two *CGI* networks with adjacency matrices $\mathbf{W}^{\mathbf{exp}}$ (experimental condition) and $\mathbf{W}^{\mathbf{cont}}$ (control), defined over the same vertex space $V$, we can obtain a differential network as $\mathbf{W}^{\mathbf{exp}} - \mathbf{W}^{\mathbf{cont}}$. Here positive values indicate interactions with higher weights in the experiment and negative values interactions with higher weights in the control. A similar operation can be performed for CCI networks. These differential networks are used for visualisation purposes (see Fig.1B of the main manuscript).

It is important to note that interactions predictions in the experiment $\mathbf{W}^{\mathbf{exp}}$ and control $\mathbf{W}^{\mathbf{cont}}$ condition should be performed on integrated and normalized single cell RNA-seq experiments to guarantee comparable weights and cell/gene spaces. An example of a valid integration and normalisation strategy using Seurat is found in our CrossTalker tutorial (based on the PMF data). In the tutorial, we used algorithms NormalizeData from the Seurat package with the following parameters: normalization.method = "LogNormalize" and scale.factor = 10000[1].

#### Implementation of Network Topological measures

When analysing a single phenotype (a single input condition), the CCI or CGI networks are used for computing the following measures (1) indegree $d_{in}$ (listener score); (2) outdegree $d_{out}$ (influencer score); (3) pagerank (importance score) $p$ and (4) betweenness centrality $b$ (mediator score) as implemented by the R package igraph[2].

When comparing two phenotypes (experiment and control), both the CGI and the CCI network are signed and directed, and there is no standard procedure to analyse such signed networks in the same way as networks with non-negative edge-weights.

In CrossTalkeR we compute the following measures from the $control$ and $exp$ networks.

$$\Delta_{in} = d_{in}(exp) - d_{in}(control) \tag{3}$$

$$\Delta_{out} = d_{out}(exp) - d_{out}(control) \tag{4}$$

$$\Delta_{betweenness} = b(exp) - b(control) \tag{5}$$

Pagerank evaluates the importance of nodes in a graph by exploring its connections, and is equal to the stationary probability distribution of a certain diffusion process on the network [3]. When applied to either the experimental condition or the control network, the pagerank of a node indicates the probability of this node to be visited by a diffusion process with random restarts[3,4] and can be used as a proxy for node importance in both CCI and CGI networks. For a given cell $c$ and state $s \in \{control, exp\}$, pagerank returns a value $\text{Pagerank}_s(c)$, which is proportional to the probability of the diffusion process to be at a cell $c$ for a given state $s$.

$$P(c|s = exp) \approx \frac{\text{Pagerank}_s(c) + \alpha}{\sum_{c\prime}(\text{Pagerank}_s(c\prime) + \alpha)} \tag{6}$$

where $\alpha$ is a regularizing constant. This is necessary as the networks do not necessarily share the same vertex set, i.e a cell or cell/gene pair is not present in a phenotype. $P(c|s = control)$ can be estimated accordingly.

We use log-odds to measure the difference in node importance with pagerank between conditions $exp$ and $control$ conditions.

$$logodds(c) = \log\left(\frac{P(c|s = exp)}{P(c|s = control)}\right). \tag{7}$$

Positive posterior log odds indicates higher probability to access a node within the experimental condition network and negative values means that there is a higher probability to access a node in the control network. We can therefore identify nodes which changes in importance in a pairs of conditions.

#### PC analysis of network properties

CrossTalkerR employs PCA as a means to combine distinct network measures for CGI networks. This analysis provides a visual way to identify features explaining distinct (and independent) properties of the network. For large CGI networks, we only indicate nodes whose values stand out, i.e. their PC1 or PC2 values are outside the 95% confidence interval assuming the PCs follow a Gaussian distribution.

See Sup. Fig. 1 for an example of the CCI network. There, we observe that megakaryocytes and Neuronal cells have increased in relevance in the experimental state: megakaryocytes as an influencer and neuronal cells by being listeners. On the other hand, other cells have decreased score in all network indices, which indicates their loss of activity/deregulation in experimental condition. A comparison of the individual phenotype networks (control and experimental condition) can also be performed via the log-odds of the importance score. While MSCs/fibroblast overall have the highest importance in individual phenotypes, we observe that megakaryocytes and neural cells have an increase in importance in the experimental condition, whereas fibroblasts a large decrease in importance due to massive loss in cellular crosstalk.

#### KEGG Annotation form Top Gene Cell pairs

To provide a functional annotation, the top $n$ ($n = 100$, as default) scored receptors/ligands for each topological measure (influencer/listener/mediator/pagerank) are used as input to the enrichKEGG function from the clusterProfiler R package. In short, we use the hypergeometric test to check if the top ranked ligand/receptors for each topological measure associated to a particular KEGG pathway is higher than expected by chance. For the contigency table, we only consider genes annotated as ligand and receptors in CellPhoneDB. We then display enriched KEGG pathways in a heatmap representation.

#### Features by CrossTalkeR and competing methods

Several computational methods have been proposed to dissect cell cell communication based on the expression of ligand and receptor genes[5]. Here we briefly revisit the methods surveyed in Armigol et. al.[5] as well as recent new LR inference methods, which work with either single cell or bulk RNA-seq as input (iTALK[6], SingleCellSignalR[7], CellPhoneDB[8], CellChat[9], ICELLNET[10], NicheNet[11], CCCExplorer[12], TalkLR[13], celltalker[14], scTensor[15] and SoptSc[16]). We are particularly interested in comparing methods regarding high level functionalities associated to ranking, visualization and comparative analysis at either the CCI or CGI network (Supplementary Table 1).

To provide a better understanding of the differences between the tools we can split the LR analysis in two main levels, low and high level. The low level is done by predicting all significant LR and respective cell type pairs given a single cell RNA seq experiment. The high level is done by exploring predicted LR interactions in terms of prioritization, data mining, dynamics and other topics of interest.

We also describe features associated to lower level tasks, i.e. tasks related to the prediction of the individual ligand-receptor interactions. These include the use of protein complex information on prediction of receptor-network pairs, the use of intra-cellular signalling information, categorization of LR pairs, as well as technical features as language and support to data containers and user friendly reports (Supplementary Table 2).

For CCI networks, several tools provide visualisation capabilities, but only few tools (CellChat[9], CrossTalkeR, iTalk[6], SingleCellSignalR[7] and TalkLR[13]) provide communication scores for cells. Of these, CellChat also explores network topology based measures such as in/out-degree, and betweenness score for single phenotype networks, but allows a phenotype comparison only in terms of the visualisation. ICELLNET[10] explores expression based values to measure condition specific cell crosstalk level.

At the Cell Gene Interaction level, several methods provide rankings for single condition networks, but only few methods address the differential analysis (CellChat[9], NicheNet[11], TalkLR[13], ICELLNET[10]). CellChat[9] builds a shared neighbour graph on pathway aggregated gene expression to find changes in two phenotypes. These are used to characterize changes in cell-gene pair activity by annotation of receptor/ligands to pathways[9]. ICELLNET, NicheNet and talkLR use cell specific expression of ligand/receptors for this. CrossTalkeR is the only tool using expression values (encoded as edge weights) and network topological measures in the CGI network for characterization of node relevance.

As described before, CrossTalkeR receives as input LR predictions from CellPhoneDB[8], which is a state-of-art method for prediction of LR interactions. It therefore includes all relevant features of this LR method considering interaction-complexes for prediction of LR pairs and the use of CellPhoneDB's complete LR database[8]. One interesting feature currently not explored in the use of intra-cellular signalling as performed by NicheNet[11]. However, this feature imposes a high computational burden in scRNA-seq data.

Altogether, CrosstalkeR is the only tool to compute network topological measures for both CCI and CGI networks, and also performs differential analysis. It has further features similar or equivalent to other methods such as visualization plots, and pathway functional analysis among others. From a technical perspective, CrossTalkeR is the only framework producing result reports using common documents format (HTML/PDF) and S4 objects, which allows for easy processing and sharing of all the data generated in each step of the framework.

| | Cell Cell Interaction Level | | | Gene Cell Interaction | | | |
|---|---|---|---|---|---|---|---|
| | **Visualization** | **Ranking** | **Differential Analysis** | **Visualization** | **Ranking** | **Differential Analysis** | **Pathway Annotation** |
| **iTALK** | Network | single | | Circos | | DE genes | |
| **SingleCellSignalR** | Circos | single | | Circos | single | DE genes | x |
| **CellPhoneDB** | | | | | single | | |
| **CellChat** | Network | single | visual | Heatmap | single & differential | statistical analysis | x |
| **IcellNet** | Network and Heatmap | | visual | DotPlot | single & differential | Visual | |
| **Nichenet** | | | | Heatmap | single & differential | | x |
| **CCCExplorer** | | | | Hypergraph | | | x |
| **CrossTalkeR** | Network | single & differential | network measures | Network & PCA | single & differential | network measures | x |
| **TalkLR** | Network | | | | single & differential | statistical measure | |
| **celltalker** | Circos | | | | single | | |
| **scTensor** | Network | single | | Network | single | | x |
| **SoptSC** | Circos/Network | | | None | single | | |

**Table 1.** Main features of ligand receptor methods regarding analysis of cell-cell and cell-gene networks

| | LR subtype | LR Complexes | Intra-signaling analysis | Language | Exportable S4 Object | Shareble Report S4 (HTML/PDF) | Built-in Database | URL |
|---|---|---|---|---|---|---|---|---|
| **iTALK** | x | | | R | | | x | https://github.com/Coolgenome/iTALK |
| **SingleCellSignalR** | | | x | R | | | x | http://www.bioconductor.org/packages/release/bioc/html/SingleCellSignalR.html |
| **CellPhoneDB** | | x | | Web and Python | | | x | https://www.cellphonedb.org/ |
| **CellChat** | | | | R | | | x | http://www.cellchat.org/ |
| **IcellNet** | x | | | R | | | x | https://github.com/soumelis-lab/ICELLNET |
| **Nichenet** | x | | x | R | | | x | https://github.com/saeyslab/nichenetr |
| **CCCExplorer** | | | x | Java | | | x | https://github.com/methodistsmab/CCCExplorer |
| **CrossTalkeR** | | x | | R | x | x | | https://github.com/CostaLab/CrossTalkeR |
| **TalkLR** | | | | R | | | x | https://github.com/yuliangwang/talklr |
| **celltalker** | | | | R | | | x | https://github.com/arc85/celltalker |
| **scTensor** | | | x | R | | | x | https://github.com/rikenbit/scTensor |
| **SoptSC** | | x | x | Matlab and R | | | | https://github.com/WangShuxiong/SoptSC |

**Table 2.** Main features of ligand receptor methods regarding the prediction of ligand-receptors.

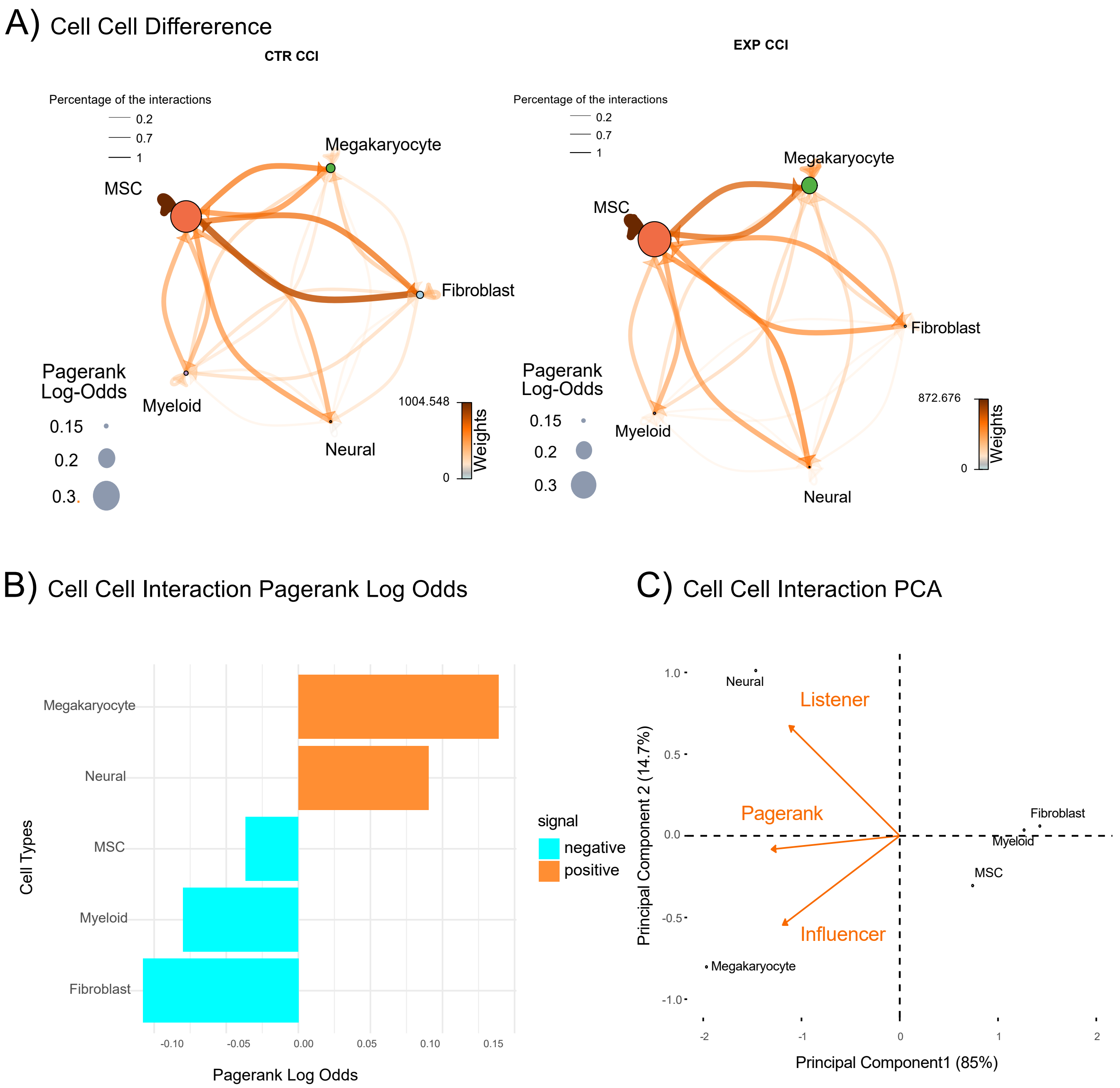


**Figure 1.** A) Cell-Cell Interaction Plot for control (control) and experimental (disease) condition in the myelofibrosis data. B) Log-odds of the importance/pagerank for networks in A. C) PCA of topological measures of the comparative CCI network. Only positive directions are shown.


**9.** Jin, S. *et al.* Inference and analysis of cell-cell communication using cellchat. *Nature communications* **12**, 1–20 (2021).

**10.** Noël, F. *et al.* Dissection of intercellular communication using the transcriptome-based framework icellnet. *Nature communications* **12**, 1–16 (2021).

**11.** Browaeys, R., Saelens, W. & Saeys, Y. Nichenet: modeling intercellular communication by linking ligands to target genes. *Nature methods* **17**, 159–162 (2020).

**12.** Choi, H. *et al.* Transcriptome analysis of individual stromal cell populations identifies stroma-tumor crosstalk in mouse lung cancer model. *Cell reports* **10**, 1187–1201 (2015).

**13.** Wang, Y. talklr uncovers ligand-receptor mediated intercellular crosstalk. *BioRxiv* (2020).

**14.** Cillo, A. celltalker (2020). URL https://arc85.github.io/celltalker/articles/celltalker.html.

**15.** Tsuyuzaki, K., Ishii, M. & Nikaido, I. Uncovering hypergraphs of cell-cell interaction from single cell rna-sequencing data. *bioRxiv* 566182 (2019).

**16.** Wang, S., Karikomi, M., MacLean, A. L. & Nie, Q. Cell lineage and communication network inference via optimization for single-cell transcriptomics. *Nucleic acids research* **47**, e66–e66 (2019).
